# The Role of Thermal Stability in AAV Titration of Engineered Variants

**DOI:** 10.1101/2024.09.11.612416

**Authors:** Emilia A Zin, Melissa Desrosiers, Tommaso Ocari, Guillaume Labernede, Camille Robert, Charlotte Izabella, Bruno Saubamea, Ulisse Ferrari, Deniz Dalkara

## Abstract

Determining the concentration of recombinant adeno-associated virus (AAV) productions, also known as titering, is crucial not only for quality control purposes but also for comparative studies of preclinical and clinical gene therapy trials. Recently, several AAVs were engineered by inserting seven amino acids at the outermost tip of the capsid’s protruding VR-VIII loop. These variants have demonstrated increased transduction capabilities over naturally occurring AAV serotypes in several studies. However, they have also been shown to produce lower yields when titered using standard techniques, raising questions about their adequacy for clinical development and use. Here, we investigated why peptide insertion onto AAV capsids reduces their titer by examining viral stocks using electron microscopy and PCR-based titering. We reveal that the DNAse digestion step, performed to eliminate free-floating DNA prior to qPCR or ddPCR, adversely impacts engineered capsid stability due to exposure to heat, artificially lowering viral titers of engineered serotypes. Titering without heating yields significantly higher titers for these variants which have melting temperatures (T_m_) close to the DNAse inactivation temperature, while titers for parental serotypes with higher T_m_ remain unchanged. Our findings provide an important new perspective for titering engineered variants with lower thermostability, especially when comparing their effectiveness to their parental serotypes.

## INTRODUCTION

Currently, adeno-associated viruses (AAVs) are the viral vector of choice for both experimental gene delivery and clinical gene therapy^1^. AAVs are small icosahedral and non-enveloped parvoviruses, around 25 nm in size^2^. There are 13 known serotypes, originally found in humans and non-human primates (NHPs), and hundreds of variants identified and developed through different methods, such as biomining, rational design and directed evolution^3^. All AAVs have the same essential structure, with 60 proteins forming the icosahedral capsid. These capsid proteins are known as VP1, VP2 and VP3 and form the capsid in a 1:1:10 ratio, respectively.

The differences in amino acid sequences of the VPs dictate the tropism of each AAV serotype or variant. For example, AAV9 shows increased tropism for the central nervous system and AAV2 is the serotype of choice for intravitreal delivery and transduction of inner retinal cells^4,5^. These differences in tropism are essential for cell targeting and increased transgene expression for therapeutic purposes. By altering or inserting specific sequences on the AAV capsid, it is possible to also alter its tropism^6,7^. Our group and others have used directed evolution to create multiple AAV vectors with enhanced tropism toward neural tissues such as the retina and the central nervous system (CNS). Of particular interest, the AAV2-7m8, AAV9-7m8 and AAV9-PHP.eB (PHP.eB) variants of AAV9 show enhanced tropism and have been widely adopted both in basic research and clinical gene therapy studies^8–10^. All of these variants were evolved by inserting seven randomized amino acids flanked by three linker amino acids. AAV2-7m8 has a 10 amino acid insertion in loop 4 of the AAV2 capsid, after amino acid position 587 whereas PHP.eB and AAV9-7m8 have amino acid insertions after position 586 of the AAV9 *Cap*. AAV2-7m8 and AAV9-7m8 provide better distribution of the transgene in retinal tissues, while PHP.eB is capable of infecting CNS neurons after intravenous delivery.

For use in gene therapies, the wild-type AAV genome is substituted by a transgene expression cassette. Genomic AAV titering using ITR (Inverted Terminal Repeat) primers and probes involves quantifying viral genome concentration through techniques like quantitative PCR (qPCR) or digital droplet PCR (ddPCR)^11,12^. Initially, the sample is treated with DNase to remove free-floating DNA, ensuring only encapsidated viral genomes are measured. Viral DNA is then extracted from the AAV particles. In qPCR, the DNA is amplified and fluorescence is measured, with the Ct values compared to a standard curve to determine the viral titer. In ddPCR, the sample is partitioned into droplets, each subjected to PCR, and the number of positive droplets is counted to directly calculate the viral genome concentration. These PCR based methods provide an indirect measure of AAV titers, essential for quality control and comparative studies in gene therapy research.

Historically, a DNase step has been included to remove all un-encapsidated viral DNA or leftover cellular DNA, to avoid contamination during the DNA amplification in the PCR reactions that follow. It is important to note that the DNase enzyme used in this step requires inactivation, which is done with a heat incubation step between 65°C to 95°C for 10 to 20 minutes, with or without the addition of EDTA^13^. Next, the capsid is opened, releasing its DNA cargo after incubation with a Proteinase K or through exposure to the PCR polymerase hot start high temperature (95°C). These titering methods have been conceived and optimized for the most frequently used viral serotypes which are thermostable within 65 to 95°C with AAV2 being of the least stable serotypes. In PBS, its melting temperature is around 66-70°C^14,15^. Other frequently used AAV serotypes, such as AAV8 and AAV9, are more stable in PBS, with melting temperatures around 72°C and 77°C respectively. Interestingly, engineered capsids such as AAV2-7m8 and AAV9-7m8 have proven to be more difficult to titer, with lower production yields when compared to parental serotypes and variable titering results. In this study, we hypothesized that the lower melting temperature of these engineered variants compared to their parental serotypes may pose a problem with the DNase inactivation step between 65°C and 95°C, which could then denature the capsids while the DNase is still active, yielding lower and variable results. Here, we show that inactivation methods that do not require an intermediate heating step result in higher and more reliable results for variants such as AAV2-7m8, AAV9-7m8 and PHP.eB.

## RESULTS

### PRODUCTION TITERS OF AAV9, AAV2, AAV9-7m8 and AAV2-7m8 SHOW DIFFERENT PROFILES OVER TIME

The viral core at the Institut de la Vision has produced thousands of AAV preparations over the years. Therefore, the historical viral titers measured in viral genomes per milliliter (vg/ml) of AAV2, AAV2-7m8, AAV9 and AAV9-7m8 can be and were compared (Figure 1A). Titers were measured with AAV2 ITR primers and qPCR after viral inactivation with traditional DNase/Proteinase K protocol, as described in Materials and Methods. AAV9 had the highest titer median, at 1.80E+14 vg/ml. AAV2 had the second-highest, at 2.23E+13 vg/ml. Meanwhile, the engineered serotypes, AAV9-7m8 and AAV2-7m8, had their median titers at 6.43E+11 and 8.79E+12 vg/ml respectively. The coefficients of variation for AAV9-7m8 and AAV2-7m8, reported in percentages, were 304% and 236% respectively, while only 187% and 91% for AAV2 and AAV9. The titer median of the engineered variants result is statistically lower than the corresponding non-engineered ones (AAV2-7m8 P=0.005, AAV9-7m8 P=1E-22).

**Figure 1.**
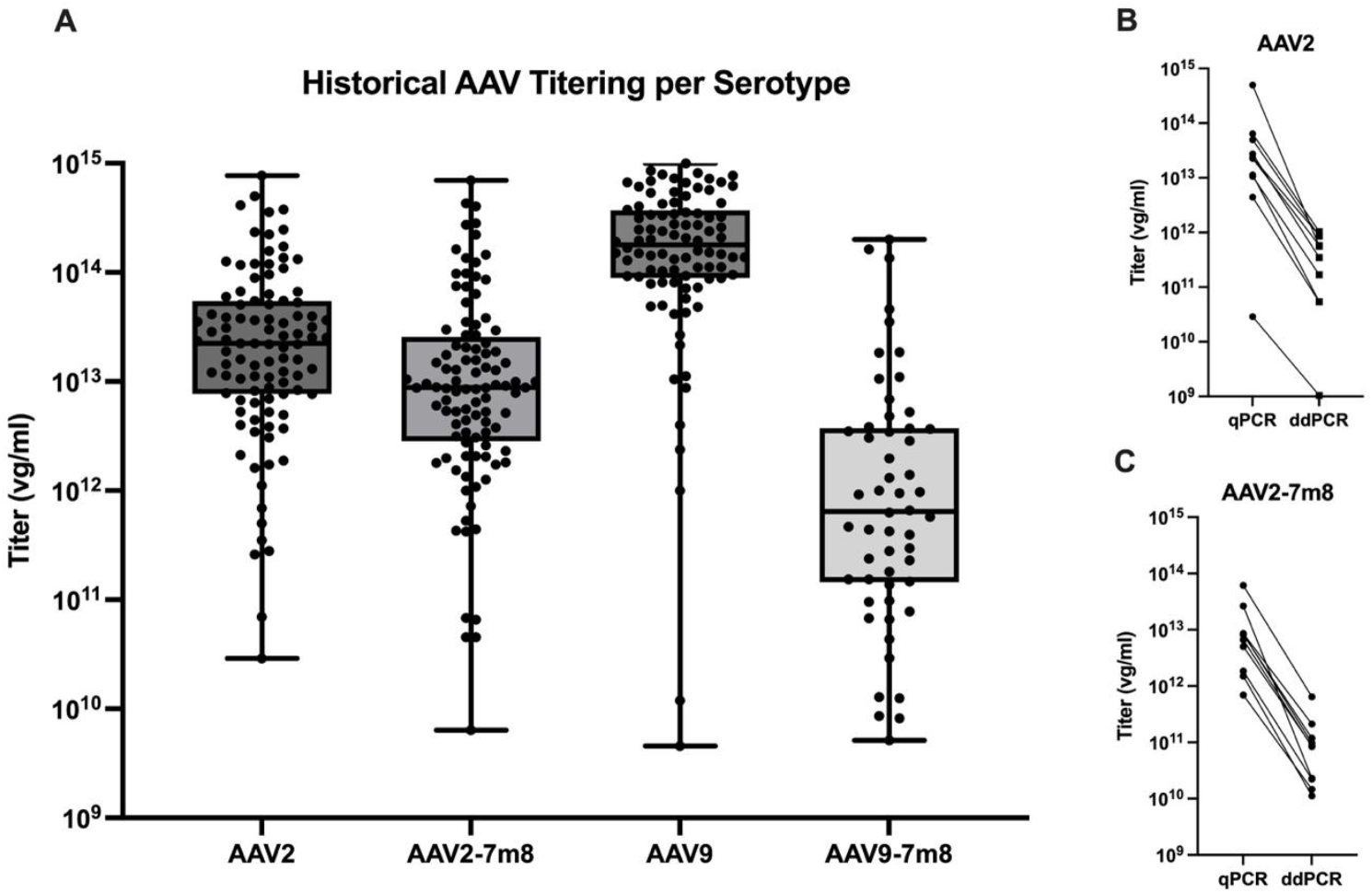
AAV titers of different productions for AAV2, AAV2-7m8, AAV9 and AAV9-7m8 over the years for the viral core at the Institut de la Vision. qPCR AAV titers by viral genomes per milliliter of production (A) post DNaseI/ProteinaseK AAV inactivation. AAV2 (B) and AAV2-7m8 (C) titers by qPCR and ddPCR. Box and whiskers plot (A), boxes show interquartile range (25% to 75%), whiskers represent maximum and minimum values, horizontal lines represent median values. AAV2 (n=99), AAV2-7m8 (n=100), AAV9 (n=92) and AAV9-7m8 (n=54), where n represents the qPCR titer value for a different AAV production. Black circles connected by lines represent the same viral production titered by either qPCR or ddPCR (B and C).

However, when comparing the historical titers obtained for AAV2 and AAV2-7m8 by qPCR and ddPCR methods, the concentration differences obtained between the former and latter methods were similar (Figure 1B and 1C). A decrease of two logs was observed for the ddPCR titer versus that obtained through qPCR. These AAV productions were packaged with different transgenes and varying cassette lengths, and only their capsid serotypes were considered for comparison.

### COMPARISON BETWEEN qPCR and ddPCR TITERS

To establish that the two-log difference between qPCR titers and ddPCR titers was consistent between different serotypes, six different capsid variants were packaged with the CAG promoter and green fluorescent protein (GFP) as the transgene in a self-complementary (sc) construct. Each cassette also contained a 21 base-pair barcode that identified the serotype in which it was packaged. These constructs were subsequently used in all experiments. Each production was inactivated with DNase/ProteinaseK and titered with both quantitative and droplet digital PCR methods–qPCR and ddPCR respectively (Figure 2). The differences between the median titer of each method ranged between 1.2 to 2.2 logs (AAV8: 1.5; AAV9: 1.8; AAV2: 1.6; PHP.eB: 1.2; AAV9-7m8: 2.2; AAV2-7m8: 1.9). While the coefficient of variation was smaller for wildtype AAVs (AAV8, AAV9 and AAV2, Figures 2A, B and C), there was a larger difference between engineered variants (PHP.eB, AAV9-7m8 and AAV2-7m8, Figures 2D, E and F).

**Figure 2.**
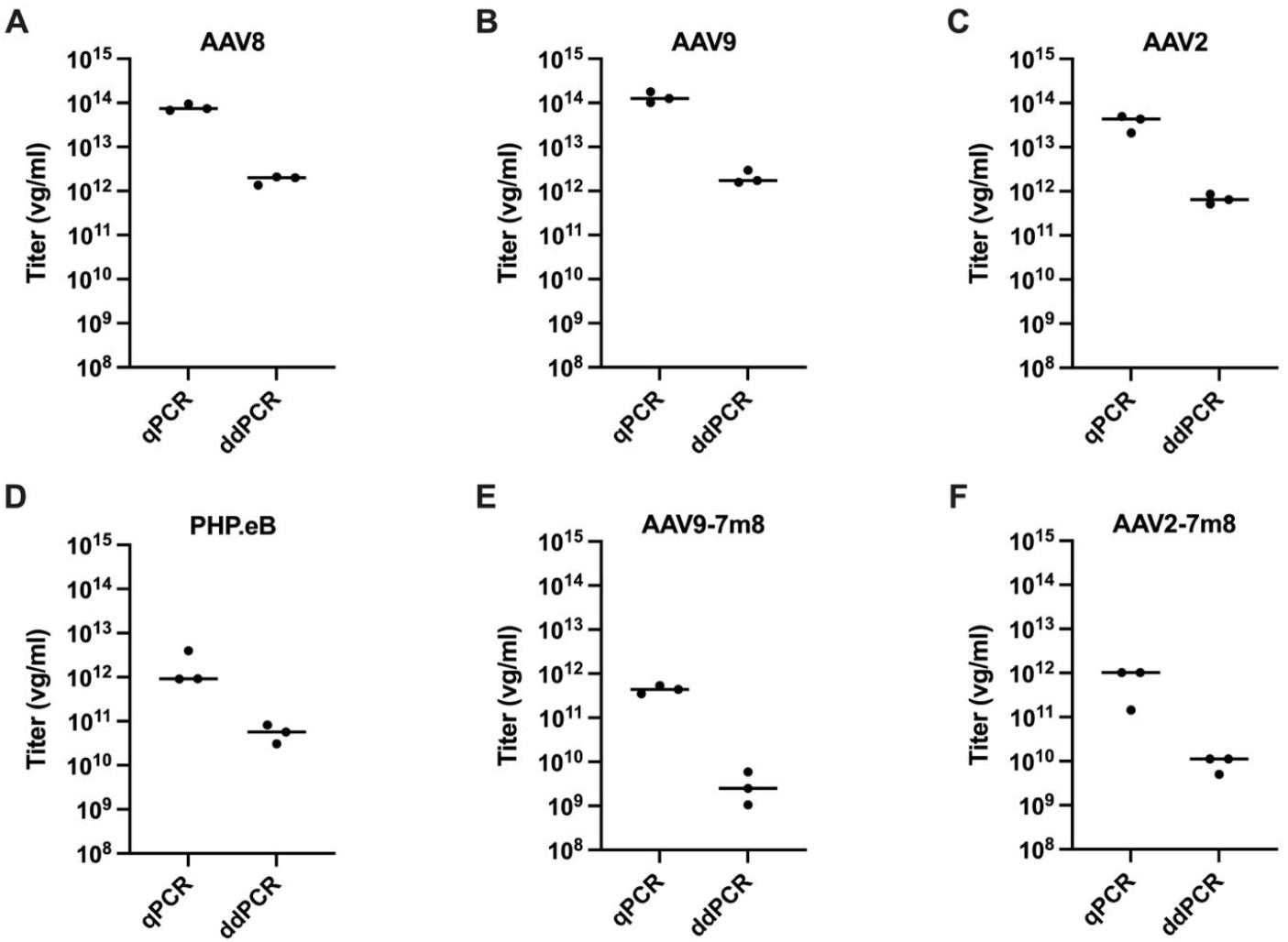
Comparison between qPCR and ddPCR titers by viral genome per milliliter (vg/ml) for individual AAV productions. AAV8 (A), AAV9 (B), AAV2 (C), PHP.eB (D), AAV9-7m8 (E) and AAV2-7m8 (F). Black circles represent different AAV productions, lines represent medians.

### THERMAL STABILITY OF ENGINEERED VARIANTS IS LOWER THAN PARENTAL SEROTYPES’

Following the observations of coefficients of variation that were larger for engineered variants than their parental serotypes, differential scanning fluorimetry (DSF) was performed to assess their capsid melting temperatures (T_m_) (Figure 3). AAV T_m_ is the point at which 50% of the viral capsid proteins are unfolded and bound with fluorescent dye. The thermal profiles for AAV8, AAV9 and AAV2 follow previously established results, where T_m_s were 73°C, 78°C and 70°C, respectively (Figure 3A and B). Meanwhile, as hypothesized, the T_m_s for engineered variants were lower than their parental serotypes. AAV9-7m8, PHP.eB and AAV2-7m8 had T_m_s of 62°C, 61°C and 66°C, respectively (Figure 3A and B). The thermal profile of the six AAV variants were then recorded using the same DSF method after a 10-minute incubation at 65°C, and results were plotted as Areas Under the Curve (AUC) (Figure 3C). There is a reduction in AUC for all AAV productions post-heat incubation, (Figure 3C).

**Figure 3.**
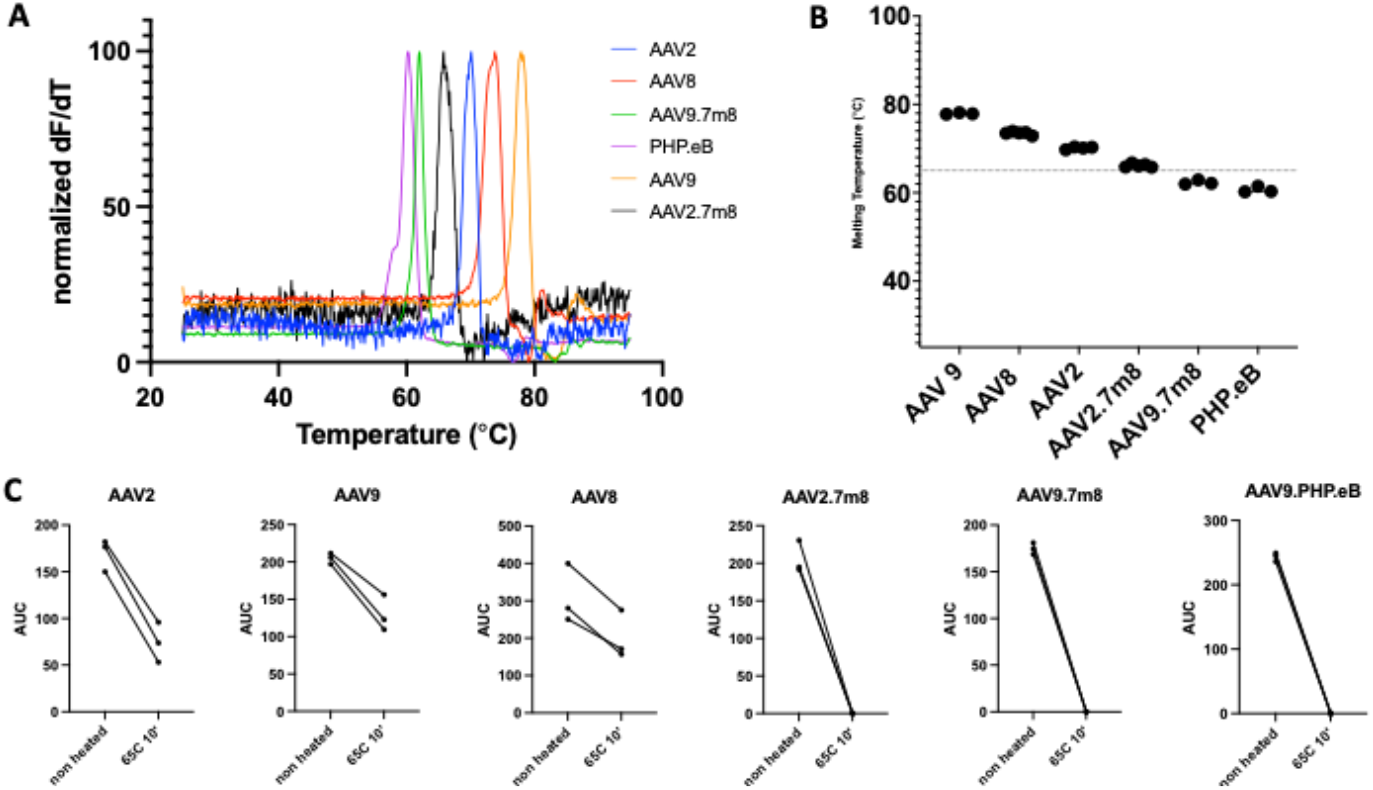
Differential scanning fluorimetry (DSF) of AAV9, AAV8, AAV2, AAV2-7m8, AAV9-7m8, PHP.eB. (A) Thermal profile (shown as normalized dF/dT, versus temperature, obtained by DSF. Example of a representative curve for each serotype.For each sample, the baseline is removed, the curve is smoothen by the second order of magnitude with 5 neighbors, and the results are normalized between 0 and 100%. (B) Plot of melting temperatures obtained from normalized dF/dT. Black circles represent melting temperature (T_m_) of individual AAV productions. Lines represent median values for each serotype.The melting temperature is determined as the temperature where the normalized derivative reaches 100%. (C) Thermal profile plotted as Area Under the Curve for each serotype, pre- and post-heat incubation at 65°C for 10 minutes. For the AUC, the dF/dT is normalized between 0 and 100% (as 100% being the maximum reached per serotype)

### ENGINEERED SEROTYPES ARE MORE SENSITIVE TO TEMPERATURE AS SEEN BY ELECTRON MICROSCOPY

After establishing that engineered variants had lower thermal stability profiles, capsids were imaged using transmission electron microscopy (TEM) to visually assess AAV capsid characteristics pre- and post-heat incubation (Figure 4). AAV8, AAV9, AAV2, AAV9-7m8, PHP.eB and AAV2-7m8 productions previously used for titering comparisons (Figure 2) or melting temperature profiles (Figure 3) were imaged directly (Figure 4A-F), or were incubated at 65°C for 10 minutes before imaging (Figure 4G-L). AAV8, AAV9 and AAV2 show a reduction in the overall number of visible AAV capsids and an increase in the number of empty capsids. Interestingly, intact or full capsids of engineered serotypes were no longer visible after heat incubation.

**Figure 4.**
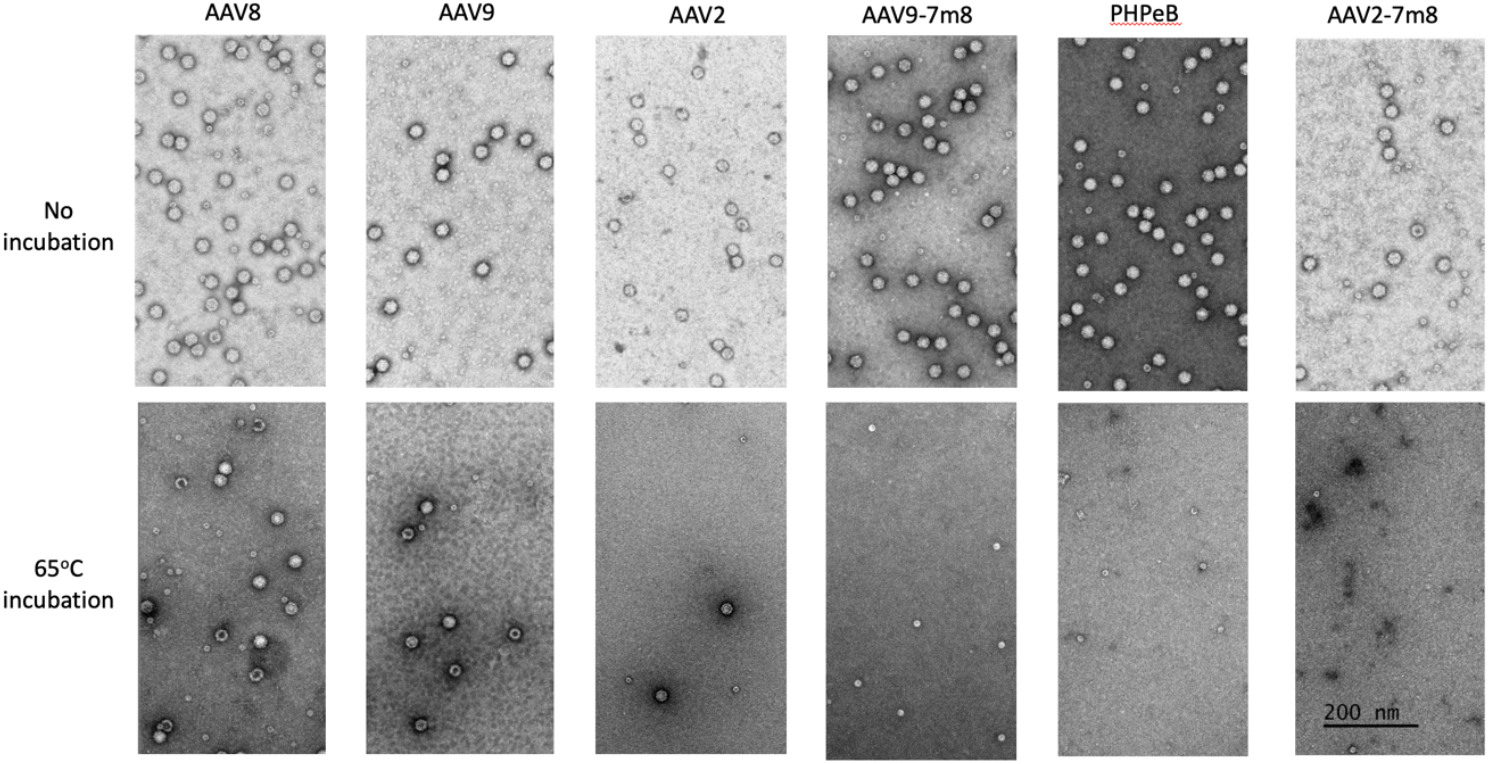
Transmission electron microscopy (TEM) images of six AAV productions with and without heat incubation. TEM images of AAV8, AAV9, AAV2, AAV9-7m8, PHP.eB and AAV2.7m8 before heat incubation (A-F, respectively) and after incubation at 65°C for 10 minutes (G-L). AAV capsids can be seen in all images except in representative images for AAV9-7m8, PHP.eB and AAV2-7m8, where only ferritin is visible (J-L). Scale bar is equal to 200 nm.

### REMOVAL OF INTERMEDIATE HEAT-INACTIVATION STEP IN TITERING PROTOCOL RESULTS IN HIGHER OVERALL TITERS

To understand if heat inactivation of DNase might indeed play a role in the final titer obtained, the same six AAV variants were titered with three different viral inactivation methods. The traditional DNase/Proteinase K method was compared to two methods that do not require heating steps. The first was Promega’s TruTiter kit, where a proprietary molecule binds to free DNA and prevents polymerases from binding in downstream reactions, thus preventing any un-encapsidated DNA from interfering in titering PCRs. The second was an inactivation protocol where DNase is only inactivated with the heat-start temperature for the ddPCR polymerase, skipping the Proteinase K step entirely, and called here the “No-Heat DNase” method. The titers for AAV8, AAV9 and AAV2 are consistent between all three methods, showing no significant differences (Figure 5A-C). However, titers obtained for the engineered variants AAV2-7m8, PHP.eB and AAV9-7m8 show a statistically significant difference between methods that do not use the heat-inactivation of DNase as an intermediary step (Figure 5D,5E and 5F). AAV2-7m8 has a median titer of 1.66E+10 vg/ml with the traditional method, while 6.72E+11 vg/ml and 1.09E+12 vg/ml with TruTiter (P=8E-4) and the DNase-only methods (P=8E-3). This represented a difference of 1.6 and 1.8 logs, respectively. PHP.eB has a median titer of 5.7E+10 vg/ml with the traditional method, while 1.23E+12 vg/ml and 2.71E+12 vg/ml with TruTiter (P=5E-3) and the DNase-only (P=1.4E-3) methods. AAV9-7m8 has a median titer of 2.51E+09 vg/ml with the traditional method, while 4.07E+11 vg/ml and 1.09E+12 vg/ml with TruTiter (P=1.4E-3) and the DNase-only (P=6E-4) methods.

**Figure 5.**
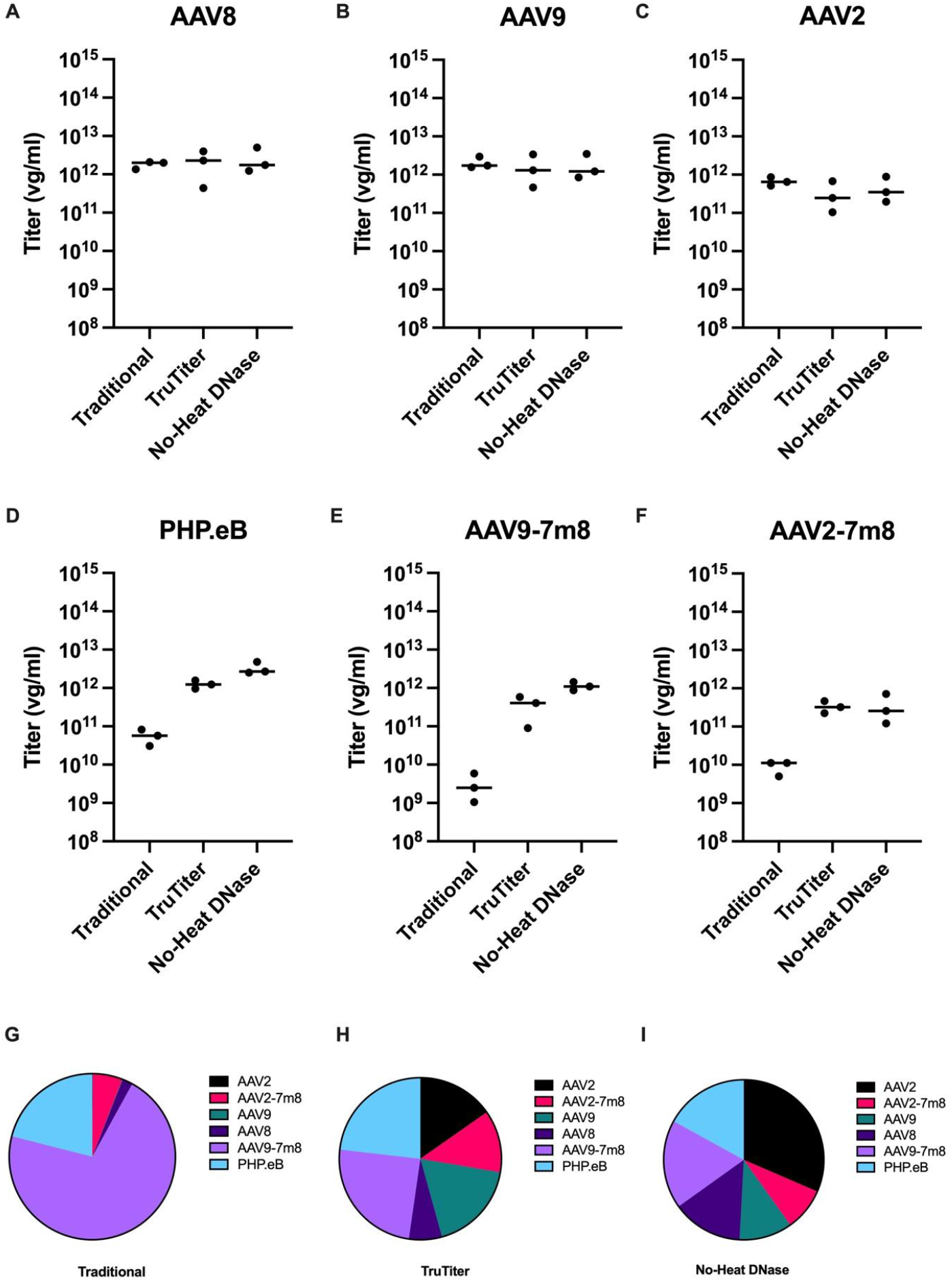
Comparison between traditional DNase/Proteinase K inactivation (Traditional), Promega’s TruTiter and protocol lacking DNase inactivation (no-heat DNase) by ddPCR quantification. AAV8 (A), AAV9 (B), AAV2 (C), PHP.eB (D), AAV9-7m8 (E) and AAV2-7m8 (F). Black circles represent different productions, lines represent medians (n=3). One-Way ANOVA with Geisser-Greenhouse’s correction performed for titration comparisons in A-F, PHP.eB P=0.048 (D), AAV9-7m8 P=0.005 (E), all other results were not significant. Equal concentration mixes were made based on these titers, and barcodes were read through next generation sequencing. Pie charts represent in percentages the resulting concentration based on titers from traditional (G), TruTiter (H) or no-heat DNase (I) inactivation.

### BARCODED AAV MIXES BASED ON TRUTITER AND NO-HEAT DNASE TITERS HAVE MORE RELIABLE CONCENTRATION RATIOS

AAV productions with a 21 bp barcode after the GFP transgene were mixed with the aim of making a solution with the same quantities of viral capsids of each serotype. The amounts were calculated based on titers obtained and reported in Figure 5A-F. The AAV mixes were then inactivated and the barcodes were amplified for NGS sequencing. As expected, TruTiter and no-heat DNase titering methods display AAV ratios that are closer to the theoretical ideal of 16.6% for each serotype. AAV variants mixed based on TruTiter results yielded 6.6 to 25.2% parts of the 100 % total (Figure 5H), while variants mixed based on the no-heat DNase method yielded 8.7 to 31.6% (Figure 5I). The traditional inactivation method yielded a mix that was highly biased towards very heat-sensitive variants, such as AAV9-7m8 and PHP.eB, which dominated the mix at 92.3% of the total (Figure 5G).

## DISCUSSION

For years, accurate, reliable and consistent AAV titration has been difficult to achieve in the gene therapy field. Although the AAV gene therapy field is advancing quickly, vector analytics and the harmonization of dosage units remain significant limitations for vector comparisons in preclinical studies and comparing results between clinical trials and even commercialization of gene therapy products. AAV reference standard materials (RSMs) have been developed as an attempt to help ensure product safety by controlling the consistency of assays used to characterize AAV stocks prepared in different laboratories but do not resolve all issues with AAV titering^16^. The most widely utilized unit of vector dosing is based on the encapsidated vector genome. Quantitative polymerase chain reaction (qPCR) is now the most common method to titer vector genomes (vg); however, significant inter- and intralaboratory variations have been documented using this technique^12^. This has been especially true with AAV2, a “finicky” capsid understood to be less stable than other natural serotypes. Moreover, with the development of engineered AAV variants, containing modifications to original AAV capsids like 7mer insertions, reliable titering became even more of a conundrum for meticulous and systematic studies of AAVs.

By quantifying the range and coefficient of variation of hundreds of AAV productions, we were able to verify quantitatively that there is indeed a larger titering variation between AAV2-7m8 and AAV9-7m8 than between their parental serotypes (Figure 1A). While this had been an anecdotal observation before, here we are able to show that when considering hundreds of different productions, the coefficient of variation for AAV9-7m8 and AAV2-7m8 was 304% and 236% respectively, while only 187% and 91% for AAV2 and AAV9. The question then became if the larger variation for engineered variants was due to titering methods themselves, potentially related to the inherent instability of the engineered AAV variants.

While the historic database of AAV titers was all obtained with qPCR, we decided for a PCR method that yielded more accurate results: ddPCR. Also, anecdotal observations indicated that this method yielded titers one or two logs lower than qPCR reactions of the same AAV production. Therefore, recent AAV productions that had been previously titered with both methods were compared. Indeed, for over 20 different productions of both AAV2 and AAV2-7m8, titers were 1.5 to 1.8 logs lower when measured by ddPCR reactions than by qPCR of the same viral inactivation (Figure 1B and C).

Therefore, we set out to test three different naturally occurring AAV serotypes, AAV8, AAV9 and AAV2, and three engineered serotypes, AAV9-7m8, PHP.eB and AAV2-7m8. AAV8 was chosen as a serotype that is frequently used for retinal gene therapy studies, as well as a serotype known to be more stable than AAV2 but less stable than AAV9^14^. AAV9 and AAV2 were chosen as the parental serotypes of the engineered AAV9-7m8, PHP.eB and AAV2-7m8. AAV9-7m8 and AAV2-7m8 are both also frequently used for retinal delivery of gene therapy cargos. Moreover, we included PHP.eB, another 7mer-insert variant based on AAV9 and popular for CNS delivery^8–10^. The six capsids (AAV8, AAV9, AAV2, AAV9-7m8, PHP.eB and AAV2-7m8) were packaged carrying the scCAG-eGFP barcoded transgene cassette for consistent comparisons.

Initially we established that the difference between qPCR and ddPCR was persistent regardless of the serotype or variant tested (Figure 2). As has been previously published, qPCR can yield an overestimation or underestimation of AAV titers due to the nature of the standard curve^17^. At the Institut de la Vision’s viral core, the standard curve is generated with linearized plasmids containing ITRs, a source that has been shown to induce an overestimation of titers.

We hypothesized that the variation in titers might come from the thermal stability of capsids. As has already been shown, AAV2 has the lowest melting temperature among the wildtype serotypes, where 50% of capsids have broken apart at 70°C in PBS. Meanwhile, AAV9 is known for its relative stability, with a melting temperature at 77°C. Considering the 65°C inactivation step of DNase during the traditional inactivation protocol of AAVs, the larger coefficient of variation for AAV2 could potentially be explained by the proximity of its T_m_ (70°C) and the 65°C inactivation temperature for DNase. Moreover, the thermal stability of engineered variants such as AAV9-7m8, PHP.eB and AAV2-7m8 had not yet been reported. Therefore, through differential scanning fluorimetry (DSF), we not only reproduced previously shown T_m_ values for wild-type serotypes AAV8, AAV9 and AAV2 (73°C, 77°C and 70°C respectively, Figure 3A & 3B), but we also showed that the T_m_ for engineered variants is lower than the parental serotypes with a striking 15°C difference for AAV9 peptide insertions (AAV9-7m8 (62°C), PHP.eB (60°C) and AAV2-7m8 (66°C)). Alarmingly, the T_m_ for all three engineered variants oscillated around the lowest possible DNase I denaturation temperature of 65°C. Consequently, we hypothesized that this incubation step was the bottleneck step that yielded lower and more variable titers for engineered variants.

To see the effects of a 65°C incubation for 10 minutes on AAV capsids, all viral productions were imaged with TEM pre- and post-incubation (Figure 4). While all AAV preparations looked as expected prior to heat incubation (Figure 4A-F), post-incubation natural serotypes had more empty capsids (Figure 4G-I). Surprisingly, all engineered capsids disappeared from TEM images, with only ferritin remaining on the EM grids (Figure 4H-J). These results corroborate the hypothesis that the 65°C incubation is damaging the AAV capsids, especially the variants that have been engineered to contain peptide insertions. To confirm these surprising results, we used the same heated sample and compared them to their non-heated counterparts with the DSF method. It was previously established that the amplitude of the curve in DSF and the amount of particles in the solution are correlated^15^. The area under the curve post heating confirmed the results seen with the electronic microscopy; a treatment of 65°C for 10 minutes reduced the number of intact particles of the parental serotype, but left no intact particles of the engineered variants (Figure 3C). It is likely that similar observations would hold true for other types of engineered variants having point mutations or loop swaps, as it has been shown that even single amino acid modifications alter thermal stability of AAV capsids.

We tested two AAV inactivation methods that did not require either DNase or DNase inactivation to prove indeed that the DNase inactivation step was the cause of variation and lower titers. First, we tested the TruTiter kit, a molecule that sequesters free-floating DNA, as well as viral DNA inside broken capsids. It does not require high temperatures for inactivation. The second method tested has already been reported, where DNase is not inactivated with heat incubation^12^. Instead, a stabilizing buffer is added to dilute the enzyme concentration and hence reduce its activity, and the preparation is taken directly to the ddPCR reaction, where the heat-start step of the polymerase at 95°C opens up the capsids and denatures the enzyme. Similarly, with TruTiter, the capsids are also opened with the heat-start step of the ddPCR polymerase.

While there were no statistically significant differences for any of the naturally occurring serotypes (Figure 5A-C), all engineered variants showed higher titers for both TruTiter and the no DNase inactivation protocol (Figure 5D-F). Moreover, these titers are more consistent with the range of natural serotype titers.

Finally, to try to corroborate these observations, we mixed all 6 AAVs according to their titers and sequenced their barcodes in order to quantify them using NGS. Massively parallel sequencing in NGS provides a more reliable and efficient method for quantifying viral genome copies compared to PCR-based approaches. NGS enables the simultaneous analysis across several samples in a single run, offering a comprehensive and unbiased measure with a broad dynamic range. This high-throughput capability allows for accurate and detailed relative quantification of barcoded genome copies, whereas PCR based methods analyze one sample at a time and can suffer from limitations such as user dependent variability. The parallel nature of NGS allowed us to objectively compare quantities of each sample mixed in hypothetical equimolar ratios. Three AAV mixes were made based on sample dilutions taking into account titers based on traditional, TruTiter or no-heat DNase titering methods. Due the unique 21 bp barcodes that corresponded to the capsid in which they were packaged, it was possible to trace the concentrations of each AAV with deep sequencing. As expected, the mix originating from the traditional titering results yielded a skewed ratio of AAVs, where the thermally unstable AAV9-7m8 and PHP.eB dominated the mix due to their underestimated titers comprising up to 92.3% of the total mix (Figure 5G). Meanwhile, the mixtures based on TruTiter and the no-heat DNase protocol showed a much more balanced profile (Figure 5H and I). These results not only show that the titers obtained with traditional methods were unreliable, but they also indicate that any experiments performed to compare the AAV infectivity profiles in such a mixture would be extremely biased. In a similar fashion, it is possible that the underestimated titers lead to misinterpretations of the effective doses of engineered AAVs in previous studies^9^.

Therefore, our study indicates that the thermal stability profile of engineered variants must be taken into consideration when titering. Engineered variants titered with traditional AAV inactivation protocols may have their titers underestimated, which has many undesirable ramifications for their downstream use. Here, we propose that traditional inactivation protocols no longer be used to titer engineered AAV variants and in studies where different AAVs need to be compared to one another. Heat inactivation is a confounding step which is best to avoid in all titering involving comparisons of AAVs with different thermostability.

## MATERIALS AND METHODS

### AAV Production

Recombinant AAV vectors were produced by the triple-transfection method as previously described^18^. AAV was purified through Iodixanol gradient and ultra-centrifugation. AAV was concentrated and resuspended in PBS with 0.01% Pluronic after buffer exchange with Amicon Ultra-50 centrifugal filter units.

### AAV Inactivation

AAV was inactivated by three different methods. The traditional and previously described protocol used DNase I (Sigma Aldrich, Missouri, USA) and Proteinase K (Sigma Aldrich). 5 µl of AAV was added to 44 µl of DNase I Buffer, followed by 1 µl of DNase I. The mixture was incubated at 37°C for 30 minutes, then heat-inactivated at 65°C for 10 minutes. 0.5 µl of Proteinase K was added to the mixture, and incubated at 50°C for 60 minutes, followed by heat-inactivation at 95°C for 20 minutes. The second inactivation method was with the TruTiter kit (Promega, Wisconsin, USA), where 5 µl of the AAV production was diluted in 45 µl of PBS-0.001% pluronic, then added to 50 µl of the diluted reaction buffer provided. The AAV is incubated for 15 minutes at 37°C, then 1 µl of the inactivation buffer is added and the solution is then incubated for 5 minutes at 20°C and 5 minutes at 95°C. Finally, the no-heat DNase inactivation method was based on a method previously published^12^. 5 µl of AAV was added to 44 µl of DNase I Buffer, followed by 1 µl of DNase I. The mixture was incubated at 37°C for 30 minutes, and 50 µl of 2x concentrated Tween20-TE buffer (Tris 10mM, EDTA 1mM, Tween20 1:10) was then added.

### AAV Titering with qPCR and ddPCR

AAV2 ITRs were quantified with 5’-GGAACCCCTAGTGATGGAGTT-3’ forward and 5’-CGGCCTCAGTGAGCGA-3’ reverse primers from Aurnhammer et al and SYBR Green (Thermo Fischer Scientific, Massachusetts, USA), using a Thermo Fisher Scientific QuantStudio 5 thermocycler and a linearized plasmid standard curve^11^. The same AAV2 ITR primers were used for ddPCR along with a FAM labeled probe 5 ’-CACTCCCTCTCTGCGCGCTCG-3 with the QX200 Bio-rad Laboratories’ droplet digital system (Bio-rad Laboratories, California, USA) and ddPCR supermix for probe (Bio-rad Laboratories, California, USA).

### Differential Scanning Fluorimetry (DSF)

The protocol was adapted from Pacouret et al^15^. 5 µl of diluted SYPRO Orange dye (S6651, Thermo Fisher Scientific, California, USA) was added to 45 µl of AAV in PBS-0.001% pluronic. The reactions were run in a Thermo Fisher Scientific QuantStudio 5 thermocycler, with experiment type as “Melt Curve” and melt curve optical settings fixed as 470 nm for excitation and 586 nm for emission. The melt curve started with two minutes at 25°C and then rose from 25°C to 95°C at 0.05°C/second. All samples were run in triplicates and averaged. Averaged data were used to calculate the derivative of the fluorescence over the derivative of the temperature (dF/dT). A peak is defined as reaching at least 20% on the Y axis and 10 adjacent points on the X axis, so the number of peaks per curve is limited to 1. The AUC is then computed by the Prism software.

### Transmission Electron Microscopy

Viruses were examined by electron microscopy after negative staining. Glow discharged Carbon/Formvar copper grids (Agar Scientific, Stansted, United Kingdom) were inverted on a drop of AAV sample for 1 min, blotted, incubated in 2% aqueous uranyl acetate for 1 min, blotted and air dried. Grids were then examined in a Jeol 1400 transmission electron microscope (Jeol, Croissy-sur-Seine, France) operated at 120 keV and equipped with a 4k x 4k RIO CMOS camera (Ametek SAS, Elancourt, France).

### Next Generation Sequencing

AAV was inactivated using the no-heat DNase method, followed by a 95°C incubation for 15 minutes. The 21 bp barcodes were amplified with 5’-GGGATCACTCTCGGCATGG-3’ forward and 5’-CTGATCAGCGAGCTCTAGGAA-3’ reverse primers and PrimeSTAR GXL DNA polymerase (Takara Bio, Shiga, Japan). They were then sequenced with NextSeq 500 1×150 (Illumina, California, USA).

### Statistical Analysis

GraphPad Prism 10.2 software (GraphPad, San Diego, CA, USA) was used to plot all graphs. Python package Scipy was used to perform all the statistical analysis. Alexander Govern test was performed in both Fig. 1 and Fig. 5. P<0.05

## ACKNOWLEDGEMENTS

This work was supported by ERC H2020 FET OPEN NEUROPA project under the grant agreement no. 863214, the Institut National de la Santé et de la Recherche Médicale (INSERM), Sorbonne Université, Agence National de Recherche (ANR) IHU FOReSIGHT (ANR18-IAHU-01), The Association UNADEV (Union Nationale des Aveugles et Déficients Visuels; National Union of the Blinds and Visually Impaired) and REGION IDF - Bourse Paris Region Fellowship CombGeneTher. EAZ is supported by the Marie Skłodowska-Curie Postdoctoral Fellowship, the project has received funding from the European Union’s Horizon Europe research and innovation programme under grant agreement No.10106380.

This work was made possible by the use of the CAPSEED, a laboratory management system dedicated to AAV production management. We would like to thank Marc Lechuga who conceived, updated and is maintaining it.

## AUTHOR CONTRIBUTIONS

Conceptualization: DD, UF, EAZ and MD; Experiments: EAZ performed the historical analysis of the viral core titers, inactivations, ddPCR, sample preparation for NGS, MD performed inactivations, ddPCR and DSF, GL and CR performed inactivations and qPCR, CI and BS performed the TEM experiments; Analysis: EAZ, MD, TO, DD and UF; Statistical Analysis: EAZ and TO; Writing (original draft): EAZ; Writing (Revision and editing): EAZ, MD, DD, UF.

## REFERENCES

1. Grimm, D. & Büning, H. Small But Increasingly Mighty: Latest Advances in AAV Vector Research, Design, and Evolution. Hum Gene Ther 28, 1075–1086 (2017).

2. Hastie, E. & Samulski, R. J. Adeno-associated virus at 50: a golden anniversary of discovery, research, and gene therapy success--a personal perspective. Human Gene Therapy 26, 257–265 (2015).

3. Srivastava, A. Rationale and strategies for the development of safe and effective optimized AAV vectors for human gene therapy. Mol. Ther. - Nucleic Acids 32, 949–959 (2023).

4. Auricchio, A. et al. Exchange of surface proteins impacts on viral vector cellular specificity and transduction characteristics: the retina as a model. Hum. Mol. Genet. 10, 3075–3081 (2001).

5. Zincarelli, C., Soltys, S., Rengo, G. & Rabinowitz, J. E. Analysis of AAV Serotypes 1–9 Mediated Gene Expression and Tropism in Mice After Systemic Injection. Mol Ther 16, 1073–1080 (2008).

6. Wu, Z. et al. Single Amino Acid Changes Can Influence Titer, Heparin Binding, and Tissue Tropism in Different Adeno-Associated Virus Serotypes. J Virol 80, 11393–11397 (2006).

7. Müller, O. J. et al. Random peptide libraries displayed on adeno-associated virus to select for targeted gene therapy vectors. Nature Biotechnology 21, 1040–1046 (2003).

8. Dalkara, D. et al. In vivo-directed evolution of a new adeno-associated virus for therapeutic outer retinal gene delivery from the vitreous. Science Translational Medicine 5, 189ra76–189ra76 (2013).

9. Khabou, H. et al. Noninvasive gene delivery to foveal cones for vision restoration. Jci Insight 3, e96029 (2018).

10. Deverman, B. E. et al. Cre-dependent selection yields AAV variants for widespread gene transfer to the adult brain. Nat Biotechnol 34, 204–209 (2016).

11. Aurnhammer, C. et al. Universal Real-Time PCR for the Detection and Quantification of Adeno-Associated Virus Serotype 2-Derived Inverted Terminal Repeat Sequences. Hum. Gene Ther., Part B: Methods 23, 18–28 (2012).

12. Wang, Y., Menon, N., Shen, S., Feschenko, M. & Bergelson, S. A qPCR Method for AAV Genome Titer with ddPCR-Level of Accuracy and Precision. Mol Ther - Methods Clin Dev 19, 341–346 (2020).

13. Green, M. R. & Sambrook, J. Removing DNA Contamination from RNA Samples by Treatment with RNase-Free DNase I. Cold Spring Harb. Protoc. 2019, pdb.prot101725 (2019).

14. Bennett, A. et al. Thermal Stability as a Determinant of AAV Serotype Identity. Mol. Ther. - Methods Clin. Dev. 6, 171–182 (2017).

15. Pacouret, S. et al. AAV-ID: A Rapid and Robust Assay for Batch-to-Batch Consistency Evaluation of AAV Preparations. Mol. Ther. 25, 1375–1386 (2017).

16. Moullier, P. & Snyder, R. O. Chapter Fifteen Recombinant Adeno-Associated Viral Vector Reference Standards. Methods Enzym. 507, 297–311 (2012).

17. Furuta-Hanawa, B., Yamaguchi, T. & Uchida, E. Two-Dimensional Droplet Digital PCR as a Tool for Titration and Integrity Evaluation of Recombinant Adeno-Associated Viral Vectors. Hum. Gene Ther. Methods 30, 127–136 (2019).

18. Choi, V. W., Asokan, A., Haberman, R. A. & Samulski, R. J. Production of Recombinant Adeno-Associated Viral Vectors. Curr Protoc Hum Genetics 53, 12.9.1-12.9.21 (2007).

